# Exploring the Binding Mechanism between Human Profilin (PFN1) and Polyproline-10 through Binding Mode Screening

**DOI:** 10.1101/418830

**Authors:** Leili Zhang, David R. Bell, Binquan Luan, Ruhong Zhou

## Abstract

The large magnitude of protein-protein interaction (PPI) pairs within the human interactome necessitates the development of predictive models and screening tools to better understand this fundamental molecular communication. However, despite enormous efforts from various groups to develop predictive techniques in the last decade, PPI complex structures are in general still very challenging to predict due to the large number of degrees of freedom. In this study, we use the binding complex of human profilin (PFN1) and polyproline-10 (P10) as a model system to examine various approaches, with the aim of going beyond normal protein docking for PPI prediction and evaluation. The potential of mean force (PMF) was first obtained from the time-consuming umbrella sampling, which confirmed that the most stable binding structure identified by the maximal PMF difference is indeed the crystallographic binding structure. Moreover, crucial residues previously identified in experimental studies, W3, H133 and S137 of PFN1, were found to form favorable hydrogen bonds with P10, suggesting a zipping process during the binding between PFN1 and P10. We then explored both regular molecular dynamics (MD) and steered molecular dynamics (SMD) simulations, seeking for better criteria of ranking the PPI prediction. Despite valuable information obtained from conventional MD simulations, neither the commonly used interaction energy between the two binding parties nor the long-term root mean square displacement (RMSD) correlates well with the PMF results. On the other hand, with a sizable collection of trajectories, we demonstrated that the average rupture work calculated from SMD simulations correlates fairly well with the PMFs (R^2^ = 0.67), making it a promising PPI screening method.

## Introduction

Polyproline recognition domains were identified in large varieties of proteins involved in intracellular signaling, including WW domains, SH3 domains, EVH1 domains, GYF domains PFN proteins, and UEV domain. ^1-7^ Their binding partners, polyproline or proline rich motifs, were found in proteins of vital importance for drug discoveries, including HIV-1 PTAP motif^7^, p53 PXXP motif^8^, Zinc finger proteins^9^, and Huntingtin protein^10^. So far, the omnipresence of polyproline is still not fully understood. For example, in Huntingtin protein, it was found that the polyglutamine length and polyproline length undergo a co-evolution from primitive organisms to human beings^10^, which may indicate a regulatory role of polyproline length in its biological functions. Yet the mechanism how Huntingtin proteins are regulated remains unknown. Therefore, it is critical to understand how proline enriched domains interact with polyproline recognition proteins, and more specifically, how the protein-protein interaction (PPI) interface conformations are determined.

PPI structure prediction is a very challenging problem computationally due to the complexity associated with the underlying multibody interactions.^11^ There are a few promising methods developed already, mostly for protein-protein docking, including ClusPro^12^, HADDOCK^13^, pyDockWeb^14^, and GRAMM-X^15^. These methods have proven successful in certain systems, especially when proteins are rigid and/or small; however, none of them could provide reliable criteria to determine the most trustworthy prediction. Every year the Critical Assessment of PRediction of Interactions (CAPRI) contest^16^ is held multiple times to test and improve various methods and protocols. Within the CAPRI framework, the contestants can submit up to 10 structures (with preferences), and the final scores are assessed based on how many hits are predicted successfully. Generally since there is no crystal structure for comparison, in past practices people used multiple models and compared the predictions, or used other empirical scoring functions to narrow down the selections. ^17-18^ Unfortunately, conflicting and confusing results from different models are frequently obtained, which complicates further assessments of the PPI structures. To overcome this shortcoming, we explore the idea of applying free energy calculations, conventional molecular dynamics (MD) simulations, and steered molecular dynamics (SMD) simulations to enhance the PPI protein docking scores (by the previously benchmarked GRAMM-X^15^) for better ranking of all predicted structures using state-of-the-art molecular modeling techniques.

Specifically, we use human profilin (PFN1) and polyproline-10 (P10) as a model system. The PPI pair PFN1-P10 is picked due to its importance and simplicity: P10 is a relatively rigid peptide adopting a polyproline II (PPII) helix^19^ that binds to surface aromatic residues of PFN1. The crystal structure of PFN1-P10 has been determined by two independent experimental groups^20-21^, proving the benchmark for the predictive computational methods. Meanwhile a challenging aspect of this system is that proline has unusual solution properties, identified as an anomalous residue in terms of its hydrophobicity ^22-23^. Thus, PFN1-P10 provides us with two possible validation tests within one system. The first is to validate if the force field guided simulations are able to reproduce experimental findings. We conducted free energy calculations to compare the crystal structure with the best structure predicted and also to investigate its associated binding mechanism. Along the latter line, we uncovered a zipping process during the binding of P10 with PFN1, highlighting the importance of W3, H133 and S137 of PFN1. The second is to examine MD simulations, steered molecular dynamics (SMD) simulations, and free energy calculations in comparison with the protein docking score (from GRAMM-X) in order to improve the ranking of docked structures and potentially go beyond protein-protein docking. We found that by following the protocol of using SMD simulation on the GRAMM-X docked structures, we were able to rank the relative stability of the PPI docked structures with reasonably high confidence.

## Methods

### Molecular docking and MD simulations

The PFN1 and P10 structures were taken from the X-ray co-crystal binding complex structure deposited in protein data bank (PDB ID: 1AWI^20^). Each of the individual protein structures was uploaded separately to the previously benchmarked protein-protein docking web server GRAMM-X^15^ (http://vakser.compbio.ku.edu/resources/gramm/grammx/) to obtain the docked complex structures (referred to as binding modes hereafter). The top 10 binding modes (ranked by scores, model to mode 10) were chosen for the latter simulations, along with the crystal structure (named modell).

MD simulations were performed with GROMACS 5.1.2 package^24^. The 11 PFN1-P10 binding modes were evaluated in simulations using the OPLS-AA force field^25^ with virtual sites for hydrogen atoms. Following similar protocols in our previous studies^26-35^, all systems were solvated in 8×8×8 nm^3^ TIP3P water boxes with 150 mM NaCl. Steepest descent method was used to minimize the solvated PFN1-P10 complexes for 10000 steps. The electrostatic interactions were calculated with particle mesh Ewald (PME) method, while the van der Waals (VDW) interactions were handled with smooth cutoffs with the cutoff distance set to 1 nm. For each mode, a 50 ps of isochoric-isothermic (NVT, 310K) simulation with 1 fs timestep were then performed to equilibrate the systems. A series of 400 ns isobaric-isothermic (NPT, 310K, 1bar) simulations with 4 fs timestep were performed during production runs.

### Interaction energy (IE)

We recorded the snapshots every 40 ps from the 400-ns MD simulations mentioned above. Water and salt were not included in the calculations of IE. With the same cutoff scheme and periodic boundary conditions as the MD simulations, we calculated the total energy of PFN1-P10 complexes (E_pFN1-p10_) and the energy of individual PFN1, P10 domains (Ep_FN1_, E_P10_) *in vacuo.* The IE is therefore calculated as:

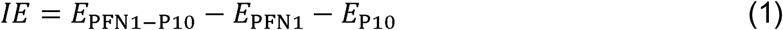

### Umbrella sampling and PMF

Umbrella sampling was performed using the minimized PFN1-P10 complexes to obtain the binding free energies. Harmonic restraints between alpha carbons (with force constants set to 1000 kJ/mol/nm^2^) were imposed on the individual domains (PFN1 and P10) to prevent them from unfolding. We utilized the center of mass (COM) distance between PFN1 and P10 as the collective variable. The window size of umbrella sampling was set as 0.1 nm, resulting in roughly 20 windows for any PFN1-P10 complexes. The simulations lasted 20 ns for each window. PME, VDW interactions and neighbor-list settings were the same as those used in MD simulations. Due to the rigidity of the system, only the translational and rotational corrections are needed to calculate the binding free energy of each binding mode.^36^ We omit this step because this correction is a constant offset for PFN1-P10 system, and absolute binding free energy is not the main interest in this study.

### Steered molecular dynamics (SMD) and rupture work

Constant velocity SMD^37^ was adopted to calculate the rupture force and rupture work of PFN1-P10 complexes. Two velocities were tested in this research (1 nm/ns and 0.1 nm/ns) along the COM distance collective variable. A total of 10 replicas were performed for each of the complexes at both velocities. Isochoric-isothermic (NVT) ensemble was used for the simulations, while all other conditions remain the same as those used in the MD simulations. The force spectra were recorded during the SMD simulations at a 0.4 ps interval. Simple maximal forces were extracted from the force spectra and the averages among 10 replicas were reported in **Table** S1. Integrations of the force spectra over the COM distances were calculated, where the maximal work in the integrated curve (see **supporting information**) was defined as the rupture work.

## Results and discussions

### GRAMM-X prefers hydrophobic binding interfaces for PFN1-P10

We use a previously benchmarked GRAMM-X^15^ to obtain the initial PFN1-P10 binding structures to determine the binding modes. The rigid docking algorithm is suitable for PFN1-P10 based on the root mean square displacements (RMSDs) of PFN1 and P10 during the MD simulations (see below). **Figure** 1 shows the top 10 docked protein-protein structures ranked by the prediction scores from mode1 to mode10 (most stable to least stable) as well as the crystal structure mode 11 (PDB ID: 1AWI) added for completeness. Note that the co-crystal structure binding mode was predicted by GRAMM-X (mode9).

**Figure 1.**
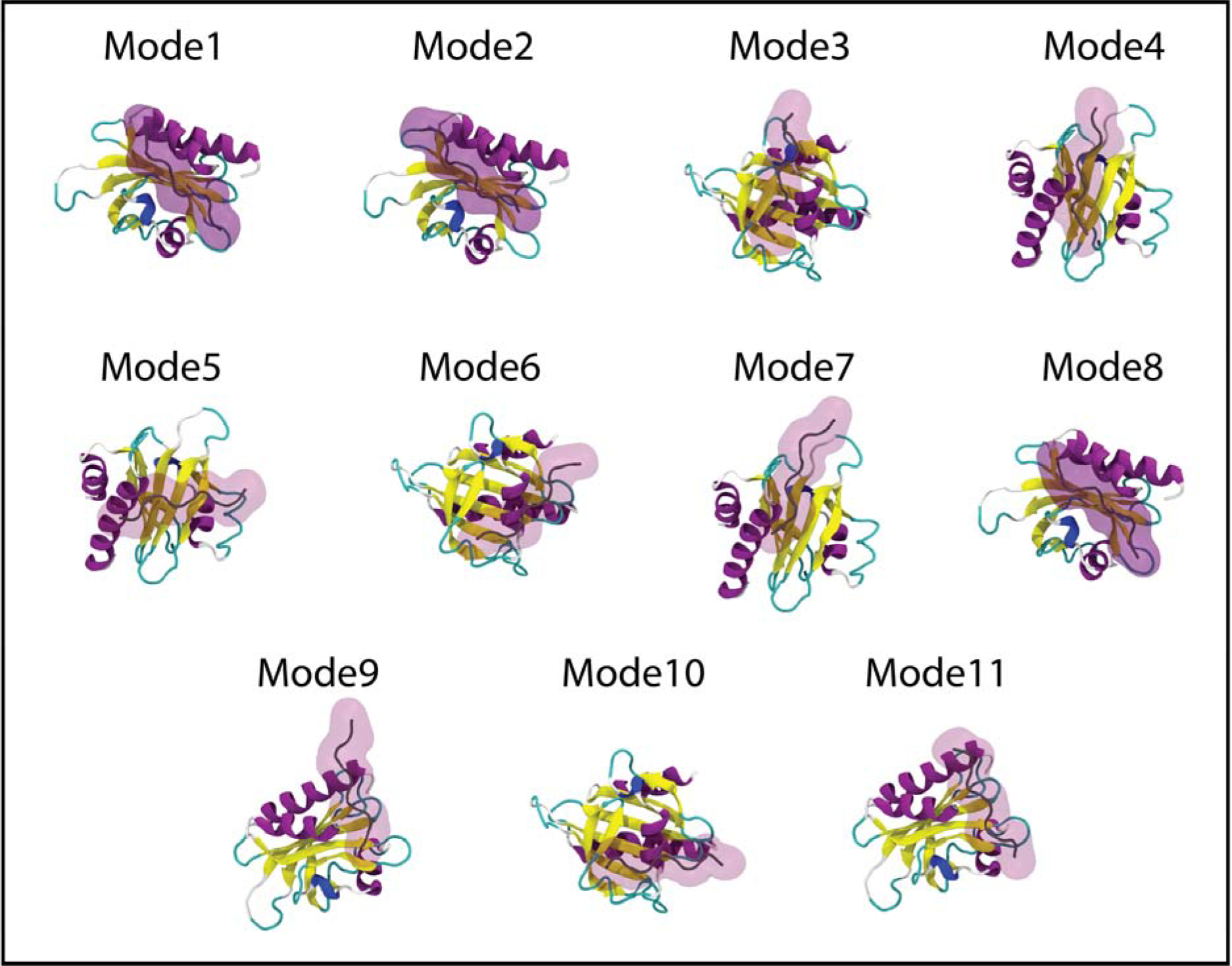
Top 10 scored human profilin (PFN1)-polyproline 10 (P10) docked structures from docking program GRAMM-X (Mode1 to Mode10). Mode11 is taken from the crystal structure (PDB ID: 1AWI).

We then analyzed interfacial residue binding for all 11 binding modes. By the proximity of the binding interfaces, we divided them into 4 categories. Category I (mode1, mode2, mode8) consists of a mostly hydrophobic binding interface on PFN1 (here we refer to PFN1 only because P10 is homogeneous and is therefore omitted later) with residues W3, N4, I7, D8, M11, A12, C16, Q17, S29, V30, W31, A32, A33, V34, P35, and K115 (62.5% hydrophobic). Category II (mode3, mode6, mode 10) utilizes a mixed hydrophobic/hydrophilic binding interface, consisting of Y24, K25, D26, S27, P28, D41, T43, P44, A45, E46, V47, G48, V49, V51, G52, K53, D54, S57, F58, N61, G62, L63, T64, and G67 (37.5% hydrophobic). Category III (mode4, mode5, mode7) is also featured by a mixed hydrophobic/hydrophilic binding interface, but consisting of a different set of residues Y59, S61, V72, I73, R74, D86, R88, A95, P96, T97, N99, E116, G117, V118, H119, G120, G121, N124, and K125 (31.6% hydrophobic). Finally, Category IV (mode9, mode11) makes the use of the binding groove from the crystal structure between the N-terminal *α* helix and C-terminal *α* helix, mainly contributed by G2, W3, N4, Y6, D26, S27, S29, W31, H133, L134, S137, and Y139 (41.7% hydrophobic). Since both the highest scores mode1 and mode2 are in Category I, GRAMM-X seems to prefer a more hydrophobic binding interface for PFN1-P10, although it also picks out mixed binding interfaces, including the co-crystal binding interface - mode9 in Category IV.

These 11 binding modes are used as the starting points for the binding free energy calculations with umbrella sampling, as well as MD simulations and SMD simulations. In the following sections we will explore these different methods in an effort to go beyond the normal (rigid) protein-protein docking for a better ranking of the binding strengths for the 11 binding modes.

### PMFs from umbrella sampling predict crystal structures to be the most stable binding modes

Umbrella sampling was adopted here to estimate the binding free energies of the 11 binding modes for the purpose of finding a practical approach of ranking binding modes from PPI docking programs and to explore the associated binding mechanism. The most rigorous way of calculating binding free energies has been discussed in multiple previous papers and is not the main focus in this study.^36, 38^ Here, we used a simplified way of finding the relative binding free energy to ensure specifically that the calculated PMF differences directly correlate with the binding mode stability, by setting the two end points to be the final binding modes (shown in **Figure** 1) and their respective unbound states (defined when the minimal distance between the two proteins, with the same conformations as in the final binding mode, is larger than 1 nm). The structures of PFN1 and P10 are restricted based on their relatively high rigidity found in the MD simulations (discussed below). This also makes the prediction of the trend of binding free energies straightforward as only a constant translational/rotational correction is needed (which is therefore omitted). Additionally, because free energy is a thermodynamic state function, the difference between the bound state and unbound state is a constant regardless of paths. With the reasons stated above, we conclude that the trend found in PMF differences between the bound state and the unbound state is the same as the trend in binding free energies.

The obtained PMF curves are shown in **Figure** 2A. The lowest binding free energies come from mode7 (Category III), mode9 and mode11 (Category IV). One important finding is that OPLS-AA force field is capable of predicting the most stable binding structure of PFN1-P10, namely the Category IV co-crystal structures (mode9, mode 11). A closer look at the binding interfaces reveals that hydrophobic interfaces such as mode1 leads to a roughly 6 kcal/mol penalty compared to the mixed binding interfaces provided by mode7, mode10 and mode11 (**Figure** 2B). Such findings further solidify the argument that polyprolines are not as hydrophobic as implied in some hydrophobicity scales^39-40^. In the next section we discuss the detailed binding mechanism unveiled from umbrella-sampling simulations.

**Figure 2.**
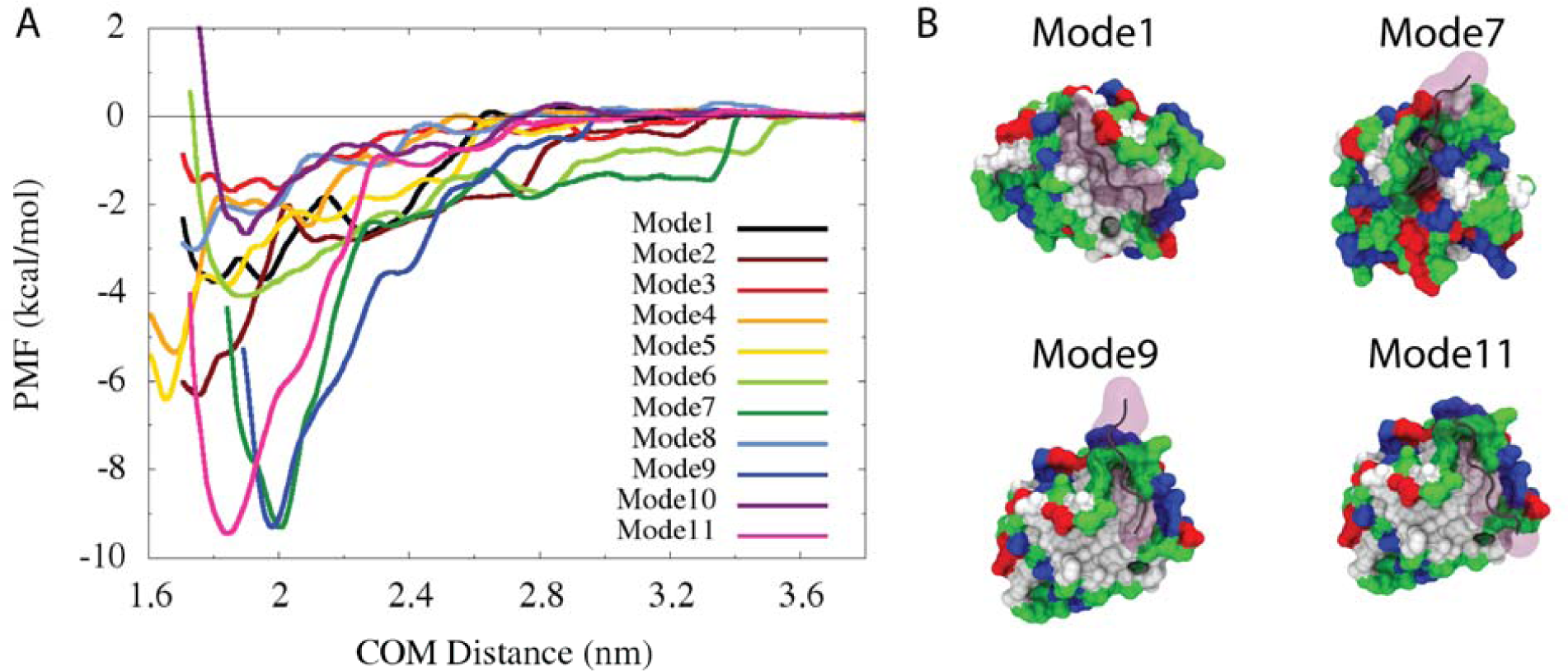
(A) PMF curves from umbrella sampling. (B) Representative binding modes where PFN1 is colored by molecular surface and residue types (hydrophobic: white; hydrophilic: green; positive: blue; and negative: red). P10 is highlighted with purple shade. The top scored mode1 mainly utilizes a hydrophobic interface. Meanwhile, the binding modes with the lowest binding free energies (mode7, mode9 and mode11) utilize a mixed hydrophobic/hydrophilic interface.

### Binding mechanism of P10 in the crystal binding pocket of PFN1

To examine the binding mechanism between PFN1 and P10, we analyzed the number of hydrogen bonds between them during the umbrella sampling simulations with Category IV binding modes (mode9, mode 11). The time-lapse counts of hydrogen bonds paired with the representative snapshots are shown in **Figure** 3 (mode9) and **Figure** S1 (mode11). Residues W3, Y6, H133, S137 and Y139 are found to contribute significantly to the PFN1-P10 contacts, in which W3^41^, H133^41^ and S137^42^ were also previously suggested to be crucial experimentally. The main contributions from tryptophans and tyrosines are consistent with previous experimental findings that prolines tend to form aromatic-proline stacking^43^, which is only seen in the crystal binding pocket (Category IV). These observations suggest that ideally, scoring functions should be tuned so that such interaction can be better captured.

**Figure 3.**
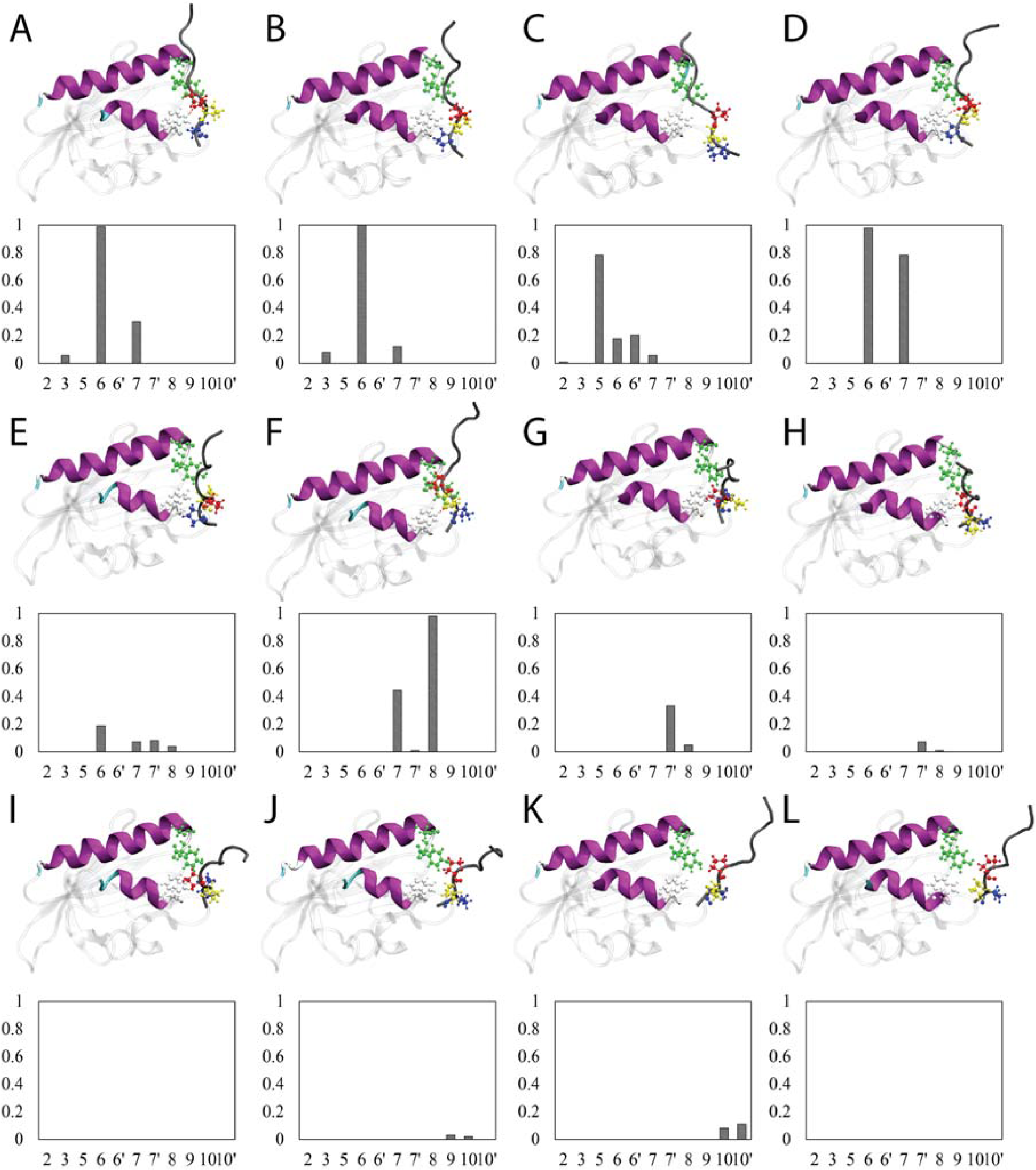
Illustrations of PFN1-P10 binding mechanism (mode9). Representative structures from window 1.6 nm to 2.7 nm are shown from (A) to (L). Only N-terminal *α* helix and C-terminal *α* helix of PFN1 are shown in visible secondary structure representations. P10 is shown as a black string representation. Typical residues involved in the interfacial hydrogen bonds are shown in bead and stick models (PFN1-W3 (white), PFN1-S137 (green), PFN1-Y139 (green), P10-P7 (red), P10-P8 (yellow) and P10-P9 (blue)). The average occurrence frequency of hydrogen bonds are plotted under each of the structures. The x axis lists all dominant hydrogen bond pairs: 2 stands for S137-P2 (the order is PFN1-P10, omitted thereafter); 3 stands for S137-P3; 5 stands for Y139-P5; 6 stands for Y139-P6; 6’ stands for W3-P6; 7 stands for W3-P7; 7’ stands for Y139-P7; 8 stands for W3-P8; 9 stands for G2-P9; 10 stands for Y139-P10; 10’ stands for S137-P10.

Next we examined how the binding process evolves when P10 approaches PFN1. A stepwise view of how hydrogen bonds form and break during umbrella sampling is shown in **Figure** 3, which illustrates how P10 binds PFN1 through a “zipping” mechanism. Hydrogen bonds are first formed between the C-terminus of P10 (residue 9∽10) and PFN1. Then the interactions gradually move along the chain to the N-terminal direction, up to the middle portion of the P10 (residue 5∽6). The aromatic-proline stacking was also examined during the binding process. We defined an aromatic-proline stacking index *(S)* based on the minimal distances from any proline side chain on P10 to the side chains of W3, Y6, Y139 from PFN1(min(*d*_p_Y6_), min(*d*_p_Y139_), respectively):

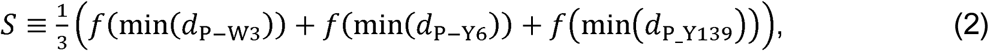

Where

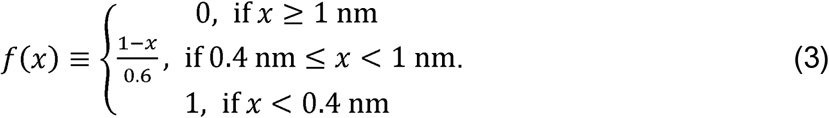

The “cutoff” switching distance of 0.4 nm was used because the average equivalent VDW radius of a residue is roughly 0.4 nm in coarse-grained models. Therefore, the close packing between two residues requires the distance to be smaller than 0.4 nm. For example, *S* = 0.8 indicates that the average minimal distance between any proline in P10 to W3, Y6, Y139 from PFN1 is 0.52 nm. **Figure** S2 shows that aromatic-proline stacking reaches the highest strength when P10 is close to PFN1 with a step function type of behavior. Combining **Figure** 3 and **Figure** S2, we observed a concerted formation of hydrogen bonds and aromatic-proline stacking when P10 approaches PFN1. This “zipping” mechanism is reproducible with two different binding modes for the crystal binding pocket: mode9 (**Figure** 3) and mode11 (**Figure** S1). Thus, in addition to the conventional aromatic-proline interaction dominated mechanism, we demonstrated that hydrogen bonds between PFN1 residues (serines, tyrosines and tryptophans) and prolines also contribute significantly to the interactions, thus resulting in a “zipping” mechanism. Such observation also helps to explain the “anomaly” of the hydrophobicity of prolines discussed before^22^.

### MD simulations disfavor hydrophobic binding interfaces

The most straightforward way of testing binding interface stability is to calculate the RMSDs of the binding modes from MD simulations. For each of the 11 initial structures prepared above, we run 400 ns of unrestrained MD simulations. We first looked at the RMSDs of individual proteins to examine their rigidity. RMSDs of PFN1 with respect to aligned initial structures are plotted in **Figure S3**A. With values always below 0.4 nm, PFN1 stays rigid throughout all simulations for all 11 binding modes, agreeing with the previous experimental findings^20^. Not surprisingly, the RMSDs of P10 with respect to the aligned initial structure (**Figure** S3B) stay below 0.3 nm. The structure of P10 stays as a polyproline II (PPII) helix throughout all of the simulations. This agrees with the widely accepted notion that polyproline should be a rigid PPII helix when the repeat length is smaller than 10.^19^ However, the RMSDs of the PFN1-P10 complex with respect to the aligned initial structure (**Figure** S3C) indicate that the binding interfaces can change significantly in some of the binding modes. For example, mode3 (bright red) reaches as high as 1.2 nm RMSD in **Figure** S3C, while PFN1 and P10 remains stable individually (**Figure** S3A and **Figure** S3B). The RMSDs of P10 with respect to the initial complex structure (not aligned, **Figure** S3D) further fortify this observation, displaying a similar trend to **Figure** S3C.

To further observe what contributes to the RMSDs, we plotted the interfacial residue frequency in **Figure** S4, with the initial occurrence frequency (0-50 ns) shown in **Figure** S4A, and final frequency (200-400 ns) shown in **Figure** S4B. The binding modes that feature major interfacial residue shifts are mode2, mode3, mode6, mode7 and mode8, mainly from Category I and Category II binding modes. More specifically, mode2 and mode8 in Category I deviate from the original hydrophobic interfacial binding, with mode2 transitioning to Category III and mode8 transitioning to Category IV, respectively. Even the relatively stable binding mode in Category I (mode1) shows a slight increase in the C-terminal region, which is the signature binding site of the co-crystal structure (Category IV). The mixed hydrophobic/hydrophilic binding interface Category II is also not very stable, with mode3 moving towards a hybrid Category II and Category III binding mode, and mode6 moving towards Category IV binding mode. Overall, all binding modes from Category I (mode1, mode2, mode8) and Category II (mode3, mode6, mode10) have tendencies to shift towards Category III and Category IV, which clearly indicates that MD simulations prefer the mixed hydrophobic/hydrophilic binding interfaces for P10. This is also phenomenologically in agreement with the umbrella sampling which predicts mode7 (Category III), mode9 and mode11 (Category IV) to be the most stable binding modes. Additionally, PFN1 C-terminus plays an important role in the final binding interfaces of mode6, mode7, mode8, mode9 and mode11. Without doubt, MD simulations (with the OPLS-AA force field) are capable to capture relevant binding features between PFN1 and P10. To further investigate if direct MD data are sufficient to warrant a reliable way of ranking the binding structures, we make several measurements discussed below.

To differentiate the short-term stability and long-term stability of the binding interfaces, we calculated the average RMSDs over 0-50 ns and 200-400 ns in **Table** S1. Interestingly, the trends of the two calculated RMSDs are vastly different. We found a positive correlation between short-term (50 ns) RMSD and PMF differences. In contrast, no correlation was found between long-term (400 ns) RMSD and PMF differences (See **Supporting Information** and **Figure** S5 for details).

Interaction energies (IEs) of PFN1-P10 are calculated with **Equation** 1 (see method for details). Similar to RMSDs, we summarized the average IEs from 0-50 ns and 200-400 ns in **Table** S1, respectively. As expected, the direct IEs are unable to recover the trend in binding free energies, regardless of short-term IEs or long-term IEs (see **Supporting Information** and **Figure** S5 for details), due to the lack of entropy contributions.

### Rank binding structures with SMD results

Based on Jarzynski’s inequality^44^, the experimental atomic force microscopy (AFM) method has proven capable of recovering the binding free energies of complicated systems such as protein-drug complexes.^45^ Concurrently, substantial efforts have been applied to adjust and optimize SMD to obtain accurate binding free energies computationally using the same inequlaity.^46-53^ Note that even though the original Jarzynski’s inequality does not require a working threshold of the rupture rate, it was largely accepted the slower the rate is, the more accurate the results will be.^47^ In general, there are limitations in using SMD for free energy calculations due to insufficient sampling sizes; however, it is still worth comparing SMD to other PPI methods. Without applying the optimization suggested for calculating binding free energies, here we test the applicability of using SMD as a fast method to rank the docked structures of PPI pairs such as PFN1-P10. We compare constant velocity SMD simulations with different pulling speeds (1 nm/ns and 0.1 nm/ns) to the properties acquired from MD simulations (RMSDs and IEs). The basic protocols of SMD can be found in the Methods section. The integrated work curves are shown in **Figure** S6, **Figure** S7. The maximal forces and rupture works are listed in **Table** S1.

With the simplest first order cumulant expansion approximation, we calculated the average rupture works from 10 replicas of SMD simulations for each of the PFN1-P10 binding modes. The rupture works calculated from two rupture speeds (1nm/ns and 0.1 nm/ns) both correlate well with the PMF differences (**Figure** 4 A, **Figure** 4 C), where R^2^ is 0.49 for 1 nm/ns SMD, and 0.67 for 0.1 nm/ns SMD. The absolute values of rupture works from 0.1nm/ns SMD (<-15 kcal/mol) are closer to the PMF differences (see **Table** S1 for details), compared to the range (−10 ∽30 kcal/mol) from 1 nm/ns SMD. This is in agreement with previous practice on free energy estimations from SMD simulations.^47^ For our purpose, instead of following the common sense of “the slower the better” in the field (0.01 or even 0.001 nm/ns), we use a relatively fast rupture speed (1 or 0.1 nm/ns) to rank the docked structures. For example, each simulation only takes ∼2.5 ns or ∼25 ns for SMD with 1 nm/ns or 0.1nm/ns rupture speed, respectively. This means that from 2.5 nsx10 replicas = 25 ns of simulations in total, a trend in PMFs can be predicted by SMD with high confidence, as compared to the 50 ns of MD simulation sampling a local well on the free energy landscape, or 20×20 = 400 ns of umbrella sampling for numerous binding modes from one PPI pair. Moreover, the replica numbers can be reduced to 6 to reach a comparable R^2^ (see **Figure** S8 A, **Figure** S8 B) which further decreases the total simulation time to 15 ns (for 1 nm/ns SMD).

**Figure 4.**
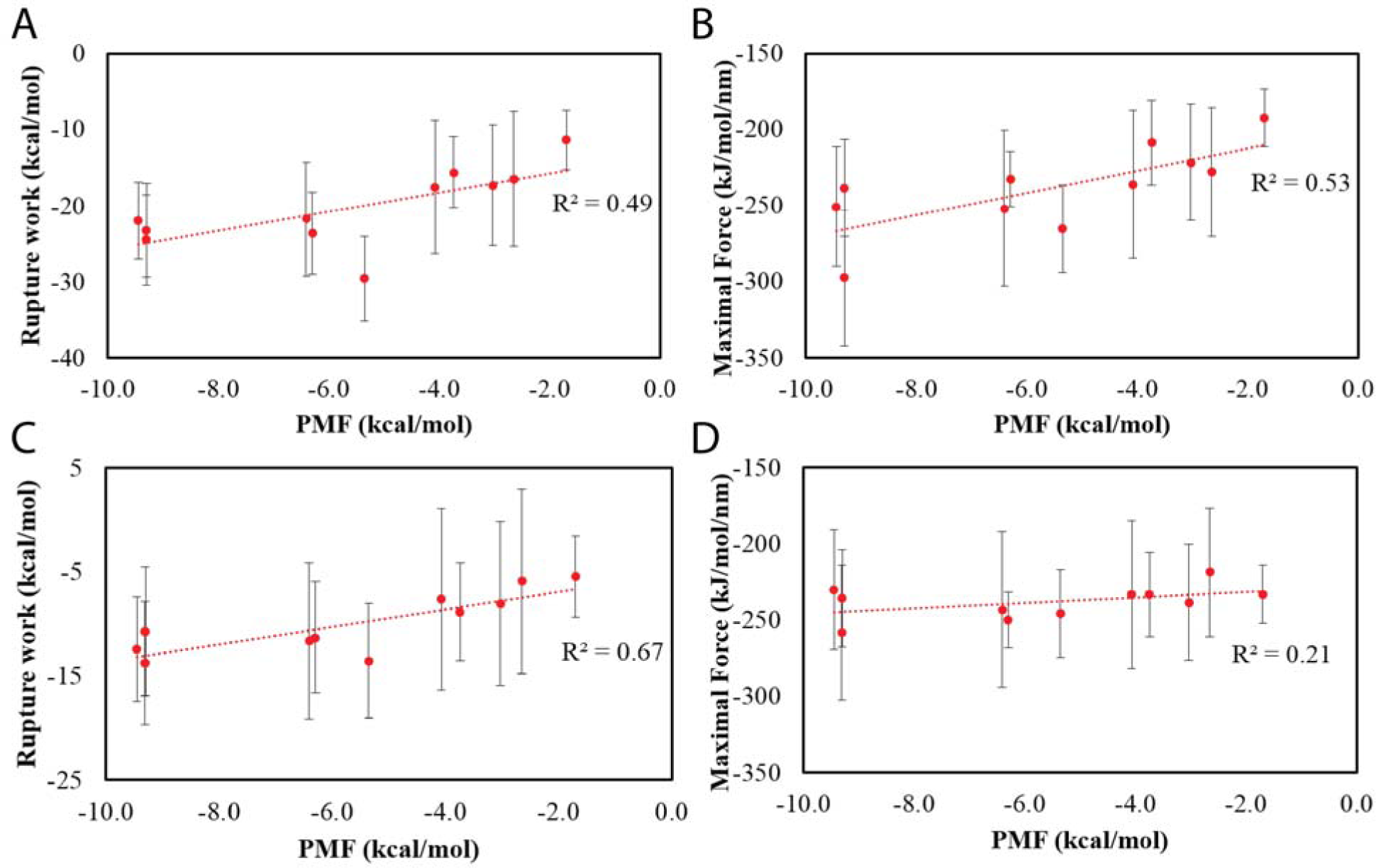
Correlation between SMD results (rupture work, maximal force) and PMF differences. (A) Rupture work calculated from 1 nm/ns SMD. (B) Maximal force recorded from 1 nm/ns SMD. (C) Rupture work calculated from 0.1 nm/ns SMD. (D) Maximal force recorded from 0.1 nm/ns SMD.

A correlation between AFM maximal forces and binding constant was reported in a cell adhesion study^54^ and an antibody-antigen unbinding study^54^. Here we investigate how maximal forces from SMD (with a much faster rupture rate) correlate with the PMFs. With two rupture rates, R^2^ is calculated to be 0.53 for 1 <u>nm</u>/ns SMD, and 0.21 for 0.1 <u>nm</u>/ns SMD, respectively. Dependent on the separation pathway selected on a free energy landscape, overall maximal forces are unsatisfactory for predicting the trend of binding free energies. The correlation may simply come from the one between maximal forces and rupture works (R^2^ is 0.65 for 1 nm/ns SMD and 0.51 for 0.1 nm/ns SMD) in this particular practice.

In **Figure** S9, we compare how the 5 properties (average RMSDs from MD over the first 50ns, maximal forces from two sets of SMD, and rupture works from two sets of SMD) rank the 11 binding modes as compared with PMFs. The correlation coefficients between the ranks are similar compared to the correlation coefficients between the absolute values (**Figure** S9 A-E), ruling out the “clustering effect” of the data (meaning the ranks may be interchangeable when absolute numbers are close). In **Figure** S9 F, we list the most stable binding modes predicted from the 5 metrics. Interestingly, mode7 is the most stable binding mode for 4 out of 5 metrics except for the rupture work (from 1 nm/ns SMD) where mode4 (in Category III along with mode7) is predicted to be the most stable. The ranks of mode11 are reliable except for maximal forces (from 0.1 nm/ns SMD). The ranks of mode9 are also high in the list but may be affected by its structural flexibility seen in the MD simulations. Notably, the top ranks derived from the rupture works from SMD correspond well to the most stable binding structures obtained from the umbrella sampling, indicating that this technique could provide a fast and reliable way of ranking the PPI binding modes from the protein docking programs such as GRAMM-X.

## Conclusion

In this paper we systematically study the PPI binding structures of PFN1-P10 using the protein docking program GRAMM-X, regular MD simulations, free energy methods (umbrella sampling) and SMD simulations, with the aim of going beyond normal protein docking for PPI prediction and evaluation. We demonstrate that the OPLS-AA force field guided umbrella sampling is able to identify the crystal structure as the most stable binding structure, which also appears in the top 10 list from GRAMM-X. Our comparative analysis shows that aromatic-proline stacking contributes the most to the stabilization of PFN1-P10 binding, along with the formation of hydrogen bonds between serines/tyrosines/tryptophans and prolines. Although regular MD simulations provide mixed information in terms of predicting the most stable binding structure, yet we find that 50 ns RMSDs might be useful with cautions for ranking the PPI prediction (see **Supporting Information** for details). On the other hand, SMD simulations provide a fast and reliable way of identifying the binding mode with the lowest binding free energy, as shown by a correlation coefficient (R^2^) as high as 0.7 (from the rank order correlation). We argue that even though the rigidity of both PFN1 and P10 may prevent us from generalizing the conclusion, by applying some constraints, SMD can be used to quickly screen the stable binding modes found from docking programs. With a clearer understanding of the advantages and disadvantages of various PPI techniques tested, we hope to expand them with a fast binding-mode-screening method to study PPI pairs found in the human interactome network^55^.

## Supporting Information

Ranking binding structures with MD results. Summarized data of MD simulations (RMSDs, IEs), umbrella sampling simulations (PMFs), and SMD simulations (maximal forces, rupture works) in **Table** S1. Binding mechanism of mode11 in **Figure** S1. Aromatic-proline stacking index measurements for umbrella sampling in **Figure** S2. RMSD of PFN, P10 and the whole complex from MD simulations in **Figure** S3. Initial binding interface and final binding interface from MD simulations in **Figure** S4. Correlation between MD simulation results and PMF differences in **Figure** S5. SMD work curves of PFN-P10 in **Figure** S6, **Figure** S7. Correlation coefficient analyses versus number of trials in **Figure** S8. Rank order correlations in **Figure** S9.

## Acknowledgement

We would like to thank Joseph A. Morrone, Hongsuk Kang, Frank Vazquez, Sangyun Lee, Serena Chen, Leticia Toledo-Sherman and Tien Huynh for their help with work. This work was partially funded by CHDI Foundation Inc., a charitable foundation that funds research into Huntington’s disease. RZ acknowledges the support of the IBM Blue Gene Science Program (W1258591, W1464125, W1464164).

